# COLOR-3D: a versatile tool for revealing novel 3D histological features

**DOI:** 10.64898/2026.05.24.725631

**Authors:** Nick King Ngai Chow, Eldric Pui Lam Tsoi, Ben Tin Yan Wong, Lichun Zhang, Tin Wai Ho, Yuqi Tan, Joshua Jing Xi Li, Hei Ming Lai

## Abstract

Hematoxylin and eosin (H&E) has been the fundamental method for visualising tissue morphology. Recent advances in tissue clearing and microscopy have enabled the observation of tissue morphology in 3D, but incomplete penetration of nucleic acid dyes has remained the bottleneck. To address this, we develop a new staining chemistry called Cyclodextrin and Organic solvent-assisted deep Labelling of ORgans in 3D (COLOR-3D), which attains the best penetration depth and homogeneity among state-of-the-art methods. We also demonstrate the scalability of COLOR-3D and its compatibility with other staining modalities. To bridge the gap between 3D histology and its wider application in biomedical research and histopathology, we develop a computational pipeline to convert 3D fluorescence images into bright-field H&E images, enabling the creation of a 3D atlas of both normal tissues and pathological specimens. Apart from qualitative observation of tissue morphology, COLOR-3D also enables quantitative analysis for studying biological phenomena. In the mouse liver, we discover rare populations of tetranuclear hepatocytes as well as m16n, t4n and t8n hepatocytes. We also propose the first structural model of the liver lobule based on 3D histology. With a more complete penetration, we reveal the following aging-related changes in tissue microarchitecture, including an increase in extreme nuclear polyploidy, and the disruption of vasculature and portal triad.

## Introduction

Hematoxylin and eosin (H&E) has been the fundamental method for visualising tissue morphology and the first-line tool in clinical pathology.^1^ Recent advances in tissue clearing and microscopy have enabled the observation of tissue morphology in 3D.^2–5^ However, the majority of biomedical research and clinical pathology still utilise 2D histology.^6–8^ The barriers to a widespread adoption of 3D histology are two-fold. First, nucleic acid dyes exhibit a limited penetration in biological tissues.^8^ Second, 3D histology requires considerable expertise and is inaccessible to most researchers and medical laboratories.^8^

Numerous methods were developed to perform tissue clearing and enhance staining depth, such as CLARITY,^9^ iDISCO^10^ and CUBIC.^11^ Representative publications that stain H&E analogues in 3D are listed in Supplementary Table 1.^12–20^ However, the penetration depth of nucleic acid dyes was still insufficient.^16^ The dye was often distributed unevenly across the depth of the tissue, which resulted in an inconsistent visualisation of tissue morphology. To improve dye penetration depth and staining homogeneity, we introduce a new staining technology called Cyclodextrin and Organic solvent-assisted deep Labelling of ORgans in 3D (COLOR-3D). COLOR-3D is a simple method that does not require specialized equipment, facilitating its adoption by clinical and research laboratories. We also validate whether COLOR-3D can be applied to different tissue types. To showcase the power of the technology, we have applied COLOR-3D to characterise the 3D histology of the mouse liver during the aging process, providing new insights in liver polyploidy and the architecture of the liver lobule.

## Results

Dye penetration can be modelled by diffusion-reaction kinetics (Equation 1).^8,16^ While the dye diffuses down a concentration gradient, dye-target binding competes with and affects diffusion. Since the outer part of the tissue depletes most of the dye, less dye is available to diffuse into the tissue’s inner part (Figure 1A). Empirically, the nucleic acid dye SYTOX Orange displayed a ‘rimming phenomenon’, where the tissue edge was brightly stained while the inner part was unstained (Figure 1B). To achieve deep and homogenous staining, we modulated diffusion-reaction kinetics by reducing dye-target binding affinity.^8^ We combined supramolecular host-guest chemistry,^21^ solvent engineering, and ionic environment optimisation. After screening for potential candidates (Supplementary Figure 1), we showed that sulfobutylether-β-cyclodextrin (SBEβCD), tetrahydrofuran (THF) and sodium chloride (NaCl) each improved penetration depth while collectively displaying a synergistic effect (Supplementary Figure 2). Unlike nucleic acid dyes, the penetration of eosin was deep and homogenous even in an aqueous solution (Figure 1B), presumably due to a lower dye-target binding affinity.

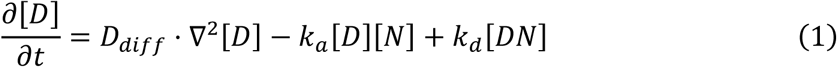

where D_diff_ denotes the diffusion coefficient; k_a_ and k_d_ denote the association and dissociation rate constants respectively; [D], [N], [DN] denote the concentrations of free dye, nucleic acid and dye-nucleic acid complex respectively.

**Figure 1:**
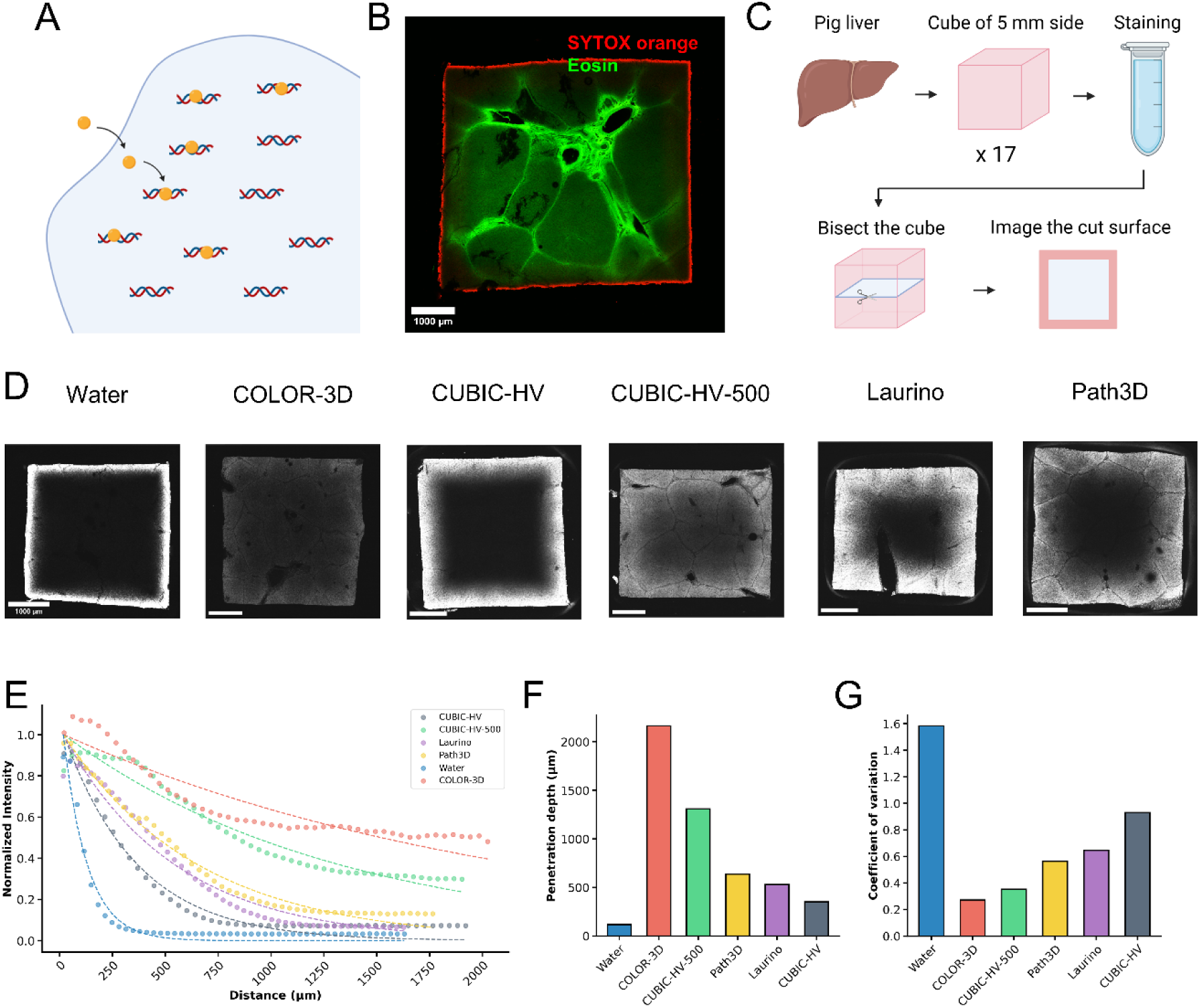
Method development process and benchmarking experiments. (A) Competition between dye-target binding and dye diffusion. (B) Rimming phenomenon of SYTOX orange, compared with deep penetration of eosin. (C) Steps for quantifying dye penetration depth. (D) Bisected surface of porcine liver. (E) Normalised staining intensity versus distance from tissue surface. The dotted line shows exponential regression. (F) Comparison of dye penetration depth. (G) Coefficient of variation of staining intensity.

We performed a head-to-head comparison with existing state-of-the-art methods (Figure 1C-D).^12,13,16^ Porcine liver tissues were cut into cubes with a side length of 5 mm, then stained with the respective protocols over a 1-day period (see Methods). We then bisected the cube and imaged the cut surface to quantify the dye penetration depth, defined as the distance from the tissue surface where staining intensity decayed by a factor of 1/e (Figure 1E). COLOR-3D attained a penetration depth of 2163 μm, which was a 17-fold increase compared to the aqueous staining solution (116 μm). COLOR-3D achieved a 65.3% increase in penetration depth over the current best method (1309 μm) (Figure 1F). Homogeneity was quantified using coefficient of variation, where a lower value indicates greater homogeneity. COLOR-3D had a coefficient of variation of 0.271, which was a 23.0% decrease over the current best method (0.353) (Figure 1G). In summary, COLOR-3D attained the best penetration depth and homogeneity among current methods, addressing a crucial bottleneck in 3D histology.

After improving nucleic acid dye penetration, we evaluated the scalability of COLOR-3D and its compatibility with other staining modalities. We stained human colon tissue measuring 8.5 mm * 8.5 mm * 3.8 mm using COLOR-3D (Figure 2A), and observed histological details including colonic glands, lymphoid aggregate, submucosa, muscularis externa, and subserosa with a blood vessel (Figure 2B). This demonstrated that COLOR-3D enabled meso-scale imaging while preserving important histological details. We also stained COLOR-3D simultaneously with other staining modalities. Antibodies and lectins showed strong staining signals (Figure 2C-D, Supplementary Figure 3), confirming their compatibility with COLOR-3D’s staining protocol. COLOR-3D also allowed multiplex staining, which was essential for mapping molecular markers across a 3D tissue (Figure 2E). For example, we simultaneously labelled the nucleus, vasculature and nervous tissues in the mouse intestine. We also labelled the nucleus, cell membrane and sinusoidal system in the mouse liver. In addition, COLOR-3D was compatible with other nucleic acid dyes including SYTO 16, YOYO 3 and DAPI, allowing additional spectroscopic combinations for multiplex staining (Figure 2F).

**Figure 2:**
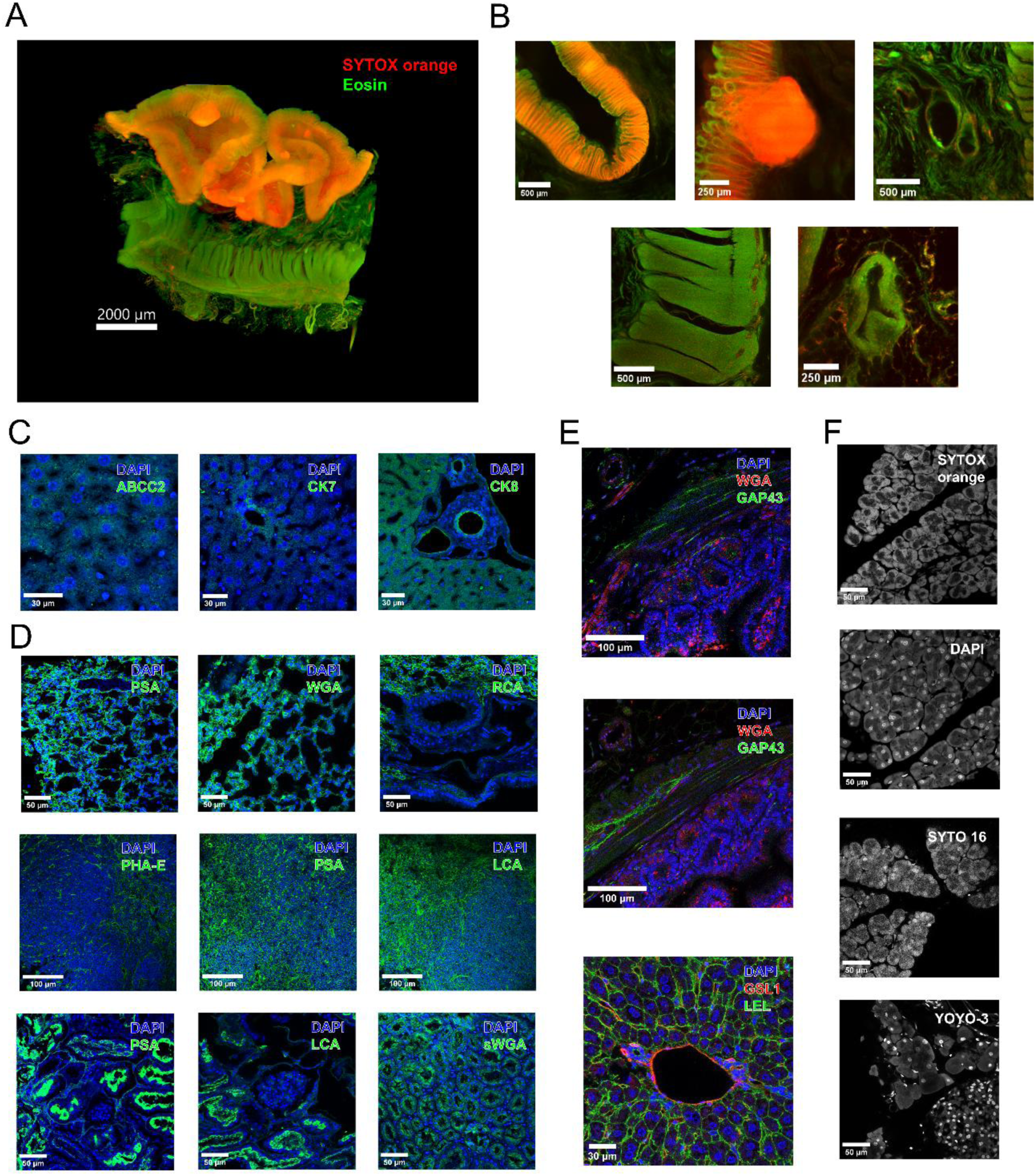
Scalability of COLOR-3D and its compatibility with other staining modalities. (A) 3D rendering of human colon tissue. (B) Enlarged views showing colonic glands, lymphoid aggregate, submucosa, muscularis externa, subserosa with a blood vessel. (C) Compatibility with immunostaining in the mouse liver. (D) Compatibility with lectin staining in the mouse lung (top row), mouse spleen (middle row) and porcine kidney (bottom row). (E) Multiplex staining. The mouse intestine shows vessels and nervous tissues. The mouse liver shows vessels, sinusoids and the cell membrane. (F) Compatibility with nucleic acid dyes.

To bridge the gap between 3D histology and its broader application in biomedical research and histopathology, it is crucial that 3D histology preserves morphological details similar to bright-field H&E images. While several existing algorithms could convert fluorescence-based images into H&E images,^13,20,22,23^ a computational pipeline that could remove imaging artifacts and adjust colour distribution was lacking. A consistent colour intensity of H&E is crucial for identifying histology structures and making pathology diagnosis. Here, we developed a computational pipeline to convert 3D fluorescence images into H&E images (Figure 3A). This multi-step, semi-automated computational pipeline comprised image preprocessing, RGB colour conversion,^13^ crosstalk removal, exponential attenuation correction, colour transfer^24^ and glitch removal. Critically, crosstalk removal and colour transfer ensured a colour distribution comparable to a real H&E image, facilitating the observation of morphological details.

**Figure 3:**
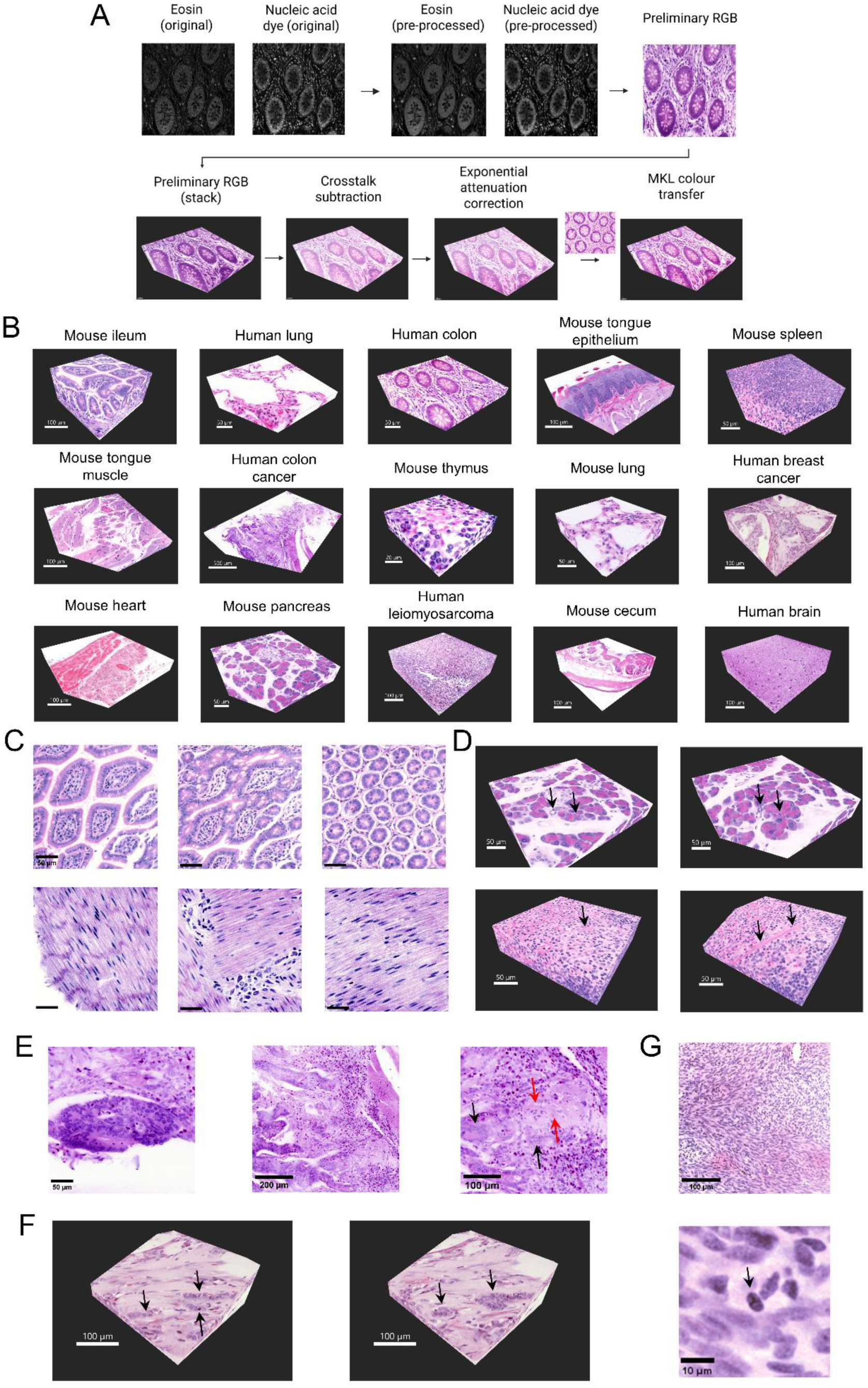
3D histology atlas created with COLOR-3D. (A) Computational pipeline to convert 3D fluorescence images into H&E images. (B) 3D images of normal tissues and pathology specimens. (C) The 3D relationship between intestinal villi, intestinal crypt and lamina propria (top row), and the myenteric ganglion embedded between mutually perpendicular smooth muscle layers (bottom row). (D) The intercalated ducts in the mouse pancreas (top row) and the splenic trabeculae in the mouse spleen (bottom row) are difficult to identify on a 2D plane (left) but revealed using the 3D image (right). (E) Human colon adenocarcinoma showing cribriform gland, submucosal invasion and perineural invasion. (F) In human breast cancer, a 2D plane may misidentify a glandular structure as a single-file pattern (left), but 3D histology can visualise the whole gland to avoid ambiguity (right). (G) Human uterus leiomyosarcoma showing haphazard smooth muscle fascicles and atypical mitosis.

We utilised COLOR-3D to create a histological atlas of normal tissues and pathology specimens. These included the gastrointestinal (GI) tract (e.g. mouse tongue, mouse ileum, mouse cecum, human colon), other internal organs (e.g. mouse lung, human lung, mouse pancreas, mouse spleen, mouse thymus, mouse heart and human brain) and human cancer tissues (e.g. colon adenocarcinoma, breast invasive ductal carcinoma, uterus leiomyosarcoma) (Figure 3B). Notably, COLOR-3D could be applied out-of-the-box with minimal optimisation or troubleshooting across multiple tissue types. Importantly, we made unique morphological observations in 3D that were difficult to view using 2D histology. For example, the 3D relationship between intestinal villi, lamina propria and intestinal glands in the mouse ileum could be viewed clearly, as well as the myenteric ganglion embedded between mutually perpendicular smooth muscle layers (Figure 3C, Supplementary Video 1).^25^ In pancreas and spleen, 3D tracing facilitated the identification of the intercalated duct and the splenic trabeculae respectively, which were difficult to identify from a single 2D cross-section (Figure 3D).^26^ In human cancer samples, we visualised 3D histological features for assessing tumour aggressiveness. In colon adenocarcinoma, we could visualise the cribriform glandular architecture, submucosal invasion and perineural invasion in a single 3D image (Figure 3E).^27^ In breast cancer, a 2D section may misidentify a glandular structure seen in invasive ductal carcinoma as a single-file pattern seen in invasive lobular carcinoma, but 3D histology could resolve this ambiguity by visualising the whole gland (Figure 3F).^28^ In the uterus, we could observe key features that differentiate a malignant leiomyosarcoma from a benign leiomyoma, such as haphazard orientation of smooth muscle fascicles and atypical mitosis (Figure 3G).^29^ We also summarised features observed in other organs (Supplementary Table 2), demonstrating that tissue morphology was well preserved in COLOR-3D-based 3D histology. To the best of our knowledge, this is the most comprehensive 3D H&E atlas available in the literature.

Apart from qualitative observation of tissue morphology, COLOR-3D also enabled quantitative analysis for studying biological phenomena, such as polyploidy in the liver. Cellular polyploidy refers to cells containing 2 or more nuclei, while nuclear polyploidy refers to a nucleus with tetraploid (4n), octoploid (8n) or 16n (Figure 4A).^30^ Liver polyploidy is of great interest due to its relationship with cellular injury, regenerative potential and carcinogenesis, but the exact proportion of polyploid hepatocytes remains debated.^31^ Existing studies used 2D histology^32^ or flow cytometry,^33^ which had respective limitations. 2D histology resulted in inaccurate quantification of the number of nuclei, nuclear size, and intensity, whereas flow cytometry only quantified nuclear intensity but not the number of nuclei (Supplementary Figure 4). We used COLOR-3D and Lycopersicon esculentum lectin (LEL) to stain and image mouse liver tissues (Figure 4B), followed by cell and nucleus segmentation with custom-trained deep learning models (Figure 4C).^34^ We segmented 142,500 hepatocytes in 4 livers and found that 47.5%, 50.4%, 1.6% and 0.4% of hepatocytes were mononuclear, binuclear, trinuclear and tetranuclear respectively (Figure 4D, Supplementary Table 3). Trinuclear hepatocytes were rarely mentioned in the literature,^35,36^ and tetranuclear hepatocytes had not been reported in a normal mouse liver previously. Then we stratified hepatocytes by cellular polyploidy and identified distinct peaks in total nuclear intensity (Figure 4E, Supplementary Figure 5). The peaks correspond to m2n, m4n, m8n and m16n for mononuclear hepatocytes, b2n, b4n and b8n for binuclear hepatocytes, t2n, t4n and t8n for trinuclear hepatocytes. The proportion of each cell type is shown in Figure 4F and Supplementary Table 3. When counted on a nuclear level, the proportions of 2n, 4n, 8n, 16n nuclei were 44.4%, 53.0%, 2.5% and 0.06% respectively (Figure 4G, Supplementary Table 4). Importantly, m16n, t4n and t8n hepatocytes were newly discovered, with examples shown in Figure 4H. We verified that these observations reflected a true biological phenomenon, but not due to image intensity artifacts or false positives in segmentation (Supplementary Figure 6). Finally, we analysed liver polyploidy using 2D sections from the 3D image. 2D sections underestimated cellular polyploidy, and distinct peaks for nuclear polyploidy could not be observed (Figure 4I). This highlights the advantage of COLOR-3D to provide an unbiased quantification of morphological parameters in 3D, whereas 2D histology may miss important features.

**Figure 4:**
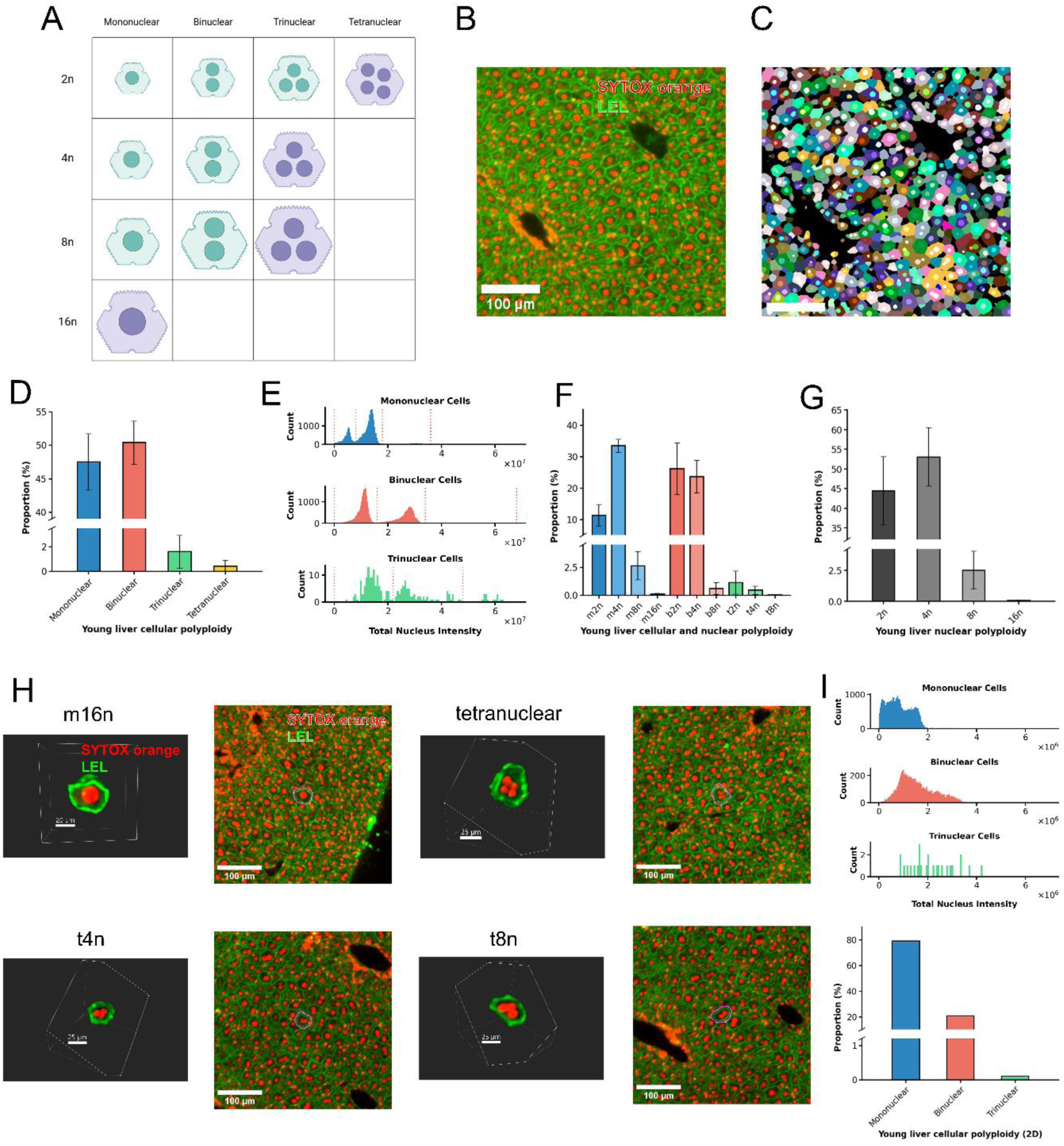
Liver polyploidy in young mice. (A) Cellular and nuclear polyploidy. Hepatocytes shown in green are currently known, while those in purple have not been reported. (B) Liver cell membrane and nuclear staining. (C) Cell and nucleus segmentation. (D) Proportion of mononuclear, binuclear, trinuclear and tetranuclear hepatocytes. (E) Distinct peaks in total nuclear intensity, stratified by cellular polyploidy status. (F) Proportion of hepatocytes with cellular and nuclear polyploidy. (G) Proportion of 2n, 4n, 8n and 16n nuclei. (H) Examples of hepatocytes with newly known polyploidy status. (I) 2D sections fail to separate the peaks in total nuclear polyploidy, while underestimating the extent of cellular polyploidy.

The advantage of COLOR-3D is further exemplified when observing multicellular, millimeter-scale structures. A liver lobule is the basic structural unit in the liver,^37^ but its 3D geometric structure remains poorly described.^38–40^ A classical description is a hexagonal prism centred on a central vein (CV) with portal veins (PV) at the vertices (Figure 5A).^25^ With COLOR-3D, we observed that both CVs and PVs are both branching structures, showing that the hexagonal prism model is oversimplified. Inspired by the 2D Voronoi diagram (Figure 5B),^41,42^ we used a modified 3D Voronoi model to represent the liver lobule. The liver lobule showed a branching pattern corresponding to the branching pattern of CV, with each lobule branch having a prismatoid shape (Figure 5C-D, Supplementary Video 2). Typically, we observed polygons of 5-7 edges if the CV is perpendicular to the plane, compared to irregular and elongated shapes if the CV is parallel to the plane (Figure 5E). While the shape of a liver lobule was irregular, the PV almost always lay on or near the lobule boundary, with a mean proximity of 85.8% (SD: 12.0%) (Figure 5F-G, Supplementary Figure 7). Although the liver lobule was studied for a century, we provided the first structural model based on 3D histology. Moreover, a liver lobule is divided into 3 zonations based on physiological functions.^43^ Here, we analysed liver polyploidy with respect to zonation (Figure 5H-I).^44^ Cellular polyploidy was concentrated in the pericentral and periportal regions, while nuclear polyploidy was concentrated in the mid-lobule region, suggesting that cellular and nuclear polyploidy have opposite biological functions. Regarding morphological parameters (Figure 5J), cell volume and nuclear volume decreased across the CV-PV axis, highlighting an important trend. Surprisingly, nuclear-cytoplasmic ratio and cell eccentricity remained constant across the CV-PV axis, revealing tight biological regulation despite the visible pleomorphism.

**Figure 5:**
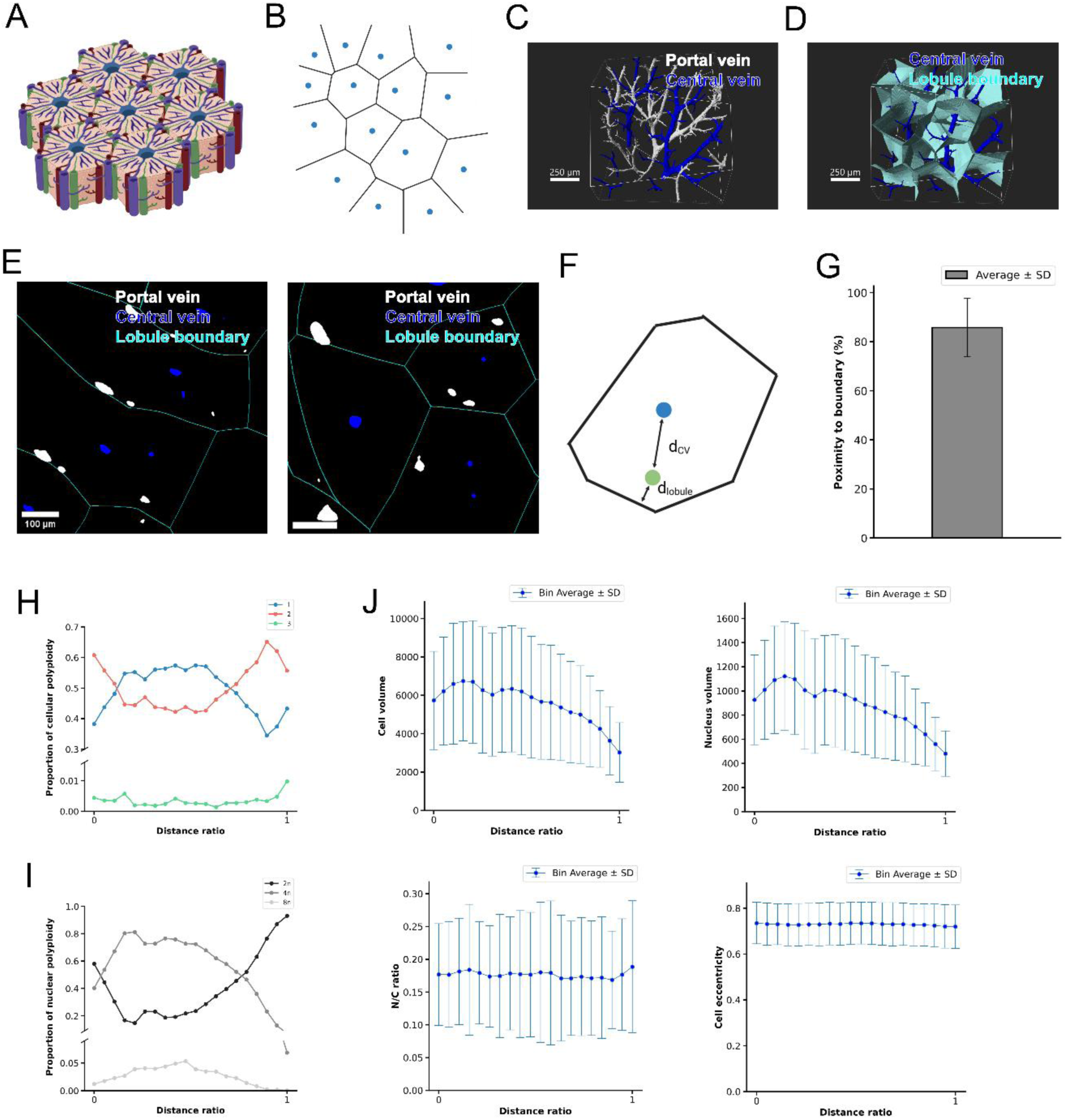
3D liver lobule and zonation analysis. (A) Traditional hexagonal prism model of the liver lobule. (B) Voronoi diagram. (C) 3D rendering of central veins (CV) and portal veins (PV). (D) 3D rendering of the liver lobule. (E) 2D sections of the liver lobule. (F) The PV is close to the lobule boundary. (G) Average proximity of the PV to the lobule boundary. (H) Cellular polyploidy across the CV-PV axis. (I) Nuclear polyploidy across the axis. (J) Regression of morphological parameters across the axis.

We also compared the histology between the young and aged mouse liver,^45–47^ identifying aging-related changes in tissue microarchitecture. We segmented 125,596 hepatocytes in 3 livers and found that 78.8%, 18.7%, 2.1% and 0.2% of hepatocytes were mononuclear, binuclear, trinuclear and tetranuclear respectively (Figure 6A-D, Supplementary Figure 5, Supplementary Table 3). When counted on a nuclear level, the proportions of 2n, 4n, 8n and 16n nuclei were 51.0%, 40.4%, 8.0% and 0.5% respectively (Figure 6E, Supplementary Table 4). Interestingly, aging was associated with a lower proportion of binuclear cells (p < 0.001), but higher proportions of extreme nuclear polyploidy including 8n (p = 0.002) and 16n (p = 0.002). This again suggests that cellular and nuclear polyploidy have opposite biological functions, where extreme nuclear polyploidy is a potential biomarker of aging and cellular injury. Moreover, we identified vascular abnormalities associated with aging. The aged liver had numerous tortuous blood vessels, in stark contrast with the straight blood vessels in the young liver (Figure 6F, Supplementary Video 3). We discovered rare phenomena, such as two separate medium-sized veins nearly touching at a single point (Figure 6G). Some veins also showed highly unusual branching patterns (Figure 6H) or even fistulae between vessel branches (Figure 6I). None of these vascular abnormalities could be visualised on a single 2D plane, highlighting the importance of 3D histology. Apart from vascular abnormalities, we also investigated the integrity of the portal triad. Previous 2D pathology studies mentioned incomplete portal triads in human liver biopsies,^48^ but their observation was limited to a 2D plane. Here, the portal triad was consistently present in the young liver but was often incomplete in the aged liver. In the aged liver, we observed a bile duct mostly present alone, but sometimes adjacent to various blood vessels (Figure 6J). These findings highlighted the extent of disruption to the liver tissue microarchitecture due to aging.

**Figure 6:**
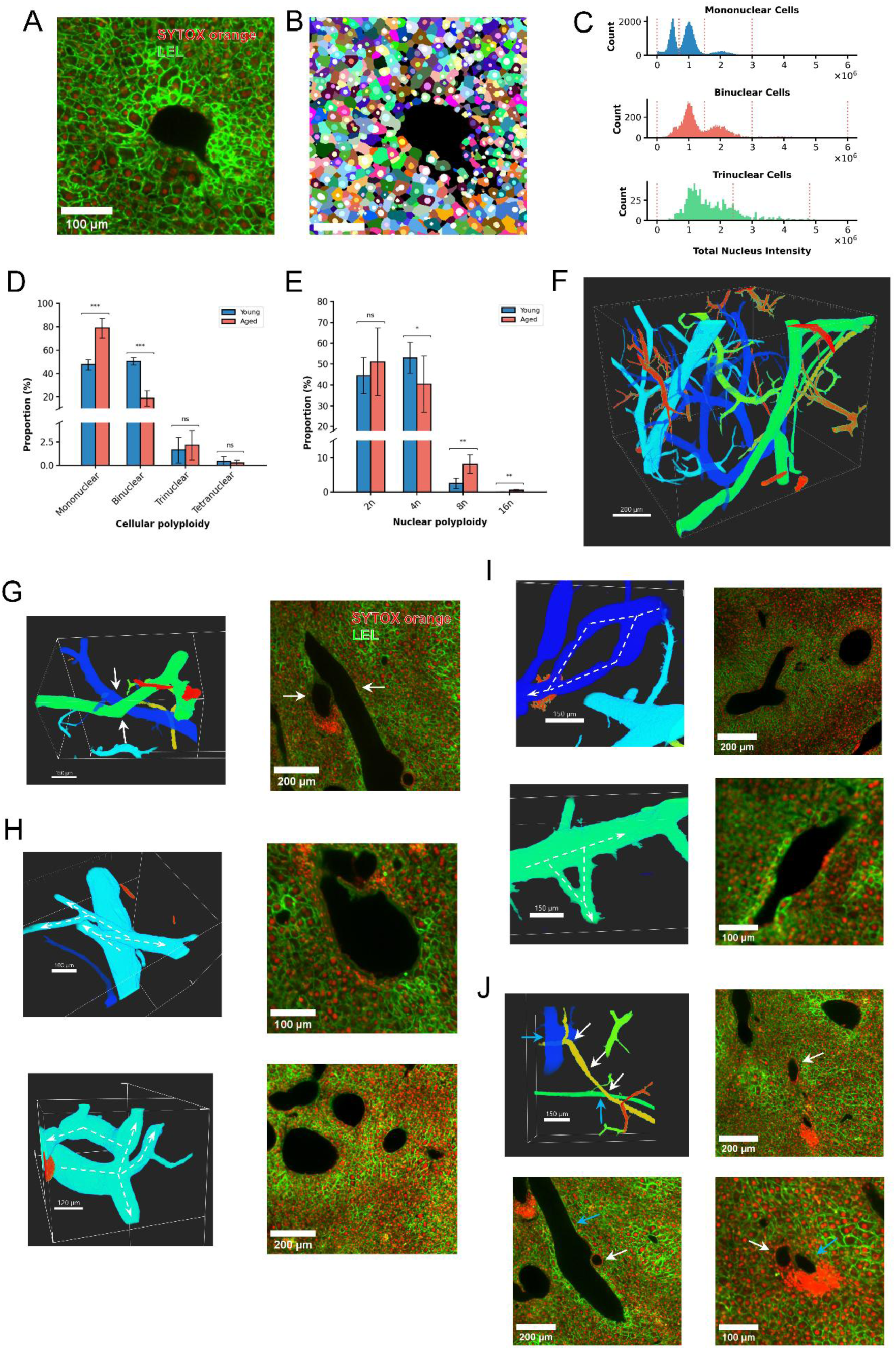
Aging-related changes in tissue microarchitecture. (A) Liver cell membrane and nuclear staining in aged mice. (B) Cell and nucleus segmentation. (C) Distinct peaks in total nuclear intensity, stratified by cellular polyploidy status. (D) Comparison of cellular polyploidy between young and aged mice. (E) Comparison of nuclear polyploidy between young and aged mice. (F) Tortuous blood vessels in the aged liver. (G) Two separate medium-sized veins nearly touching at a single point. (H) Unusual vessel branching patterns. (I) Fistulae between vessel branches. (J) The bile duct is mostly present alone (top right), but sometimes adjacent to various blood vessels (bottom row).

## Discussion

We have developed COLOR-3D, a novel staining technology which improves nucleic acid dye penetration for 3D histology. Through combining supramolecular host, organic solvent and ionic environment, COLOR-3D achieves the best dye penetration depth and homogeneity among state-of-the-art methods. Penetration depth and homogeneity is crucial for visualising morphological details and for downstream quantitative analysis.^8,16,49^ Moreover, we have validated COLOR-3D extensively, such as its ability to stain and image full-thickness human colon tissues. COLOR-3D is also highly versatile, showing compatibility with immunostaining, lectin staining, and the use of other nucleic acid dyes.

Here, by extending H&E into 3D, we aim to revolutionise how tissue morphology is studied. Our integrated colour conversion pipeline serves as the basis for a 3D histology atlas, which includes the GI tract, other internal organs, and human cancer tissues. We have identified interesting 3D morphological features which serve as a starting point for researchers and histopathologists to study 3D histology. More importantly, COLOR-3D does not require specialised equipment and is ready for immediate use across multiple tissue types, therefore expanding its application in biomedical research and clinical pathology.

In addition, we have shown that COLOR-3D is not limited to qualitative observation, but also allows an in-depth, quantitative analysis of liver histology. While previous studies have quantified polyploidy in the mouse liver,^32,33,35,36^ here we characterise the true extent of cellular and nuclear polyploidy using 3D histology. We discover rare populations of tetranuclear, m16n, t4n and t8n hepatocytes, which were overlooked by previous studies. The aged liver shows differences in polyploidy status, suggesting polyploidy as a biomarker for aging and cellular injury. We have also analysed cellular morphology parameters with respect to zonation, revealing tight biological regulation despite the visible pleomorphism. Furthermore, we define the 3D shape of the liver lobule by combining innovations in both staining and mathematical analysis, challenging the traditional hexagonal prism model.^25,26^ The aged liver shows significant disruption in vasculature and liver lobule organisation, which is another potential biomarker for aging and various pathological states. This highlights COLOR-3D as a valuable tool to solve complex histology puzzles and make novel biological discoveries.

Nevertheless, we acknowledge several limitations in our study. First, although COLOR-3D improves dye penetration depth and staining homogeneity, the maximum size of the tissue remains limited to imaging constraints, such as light attenuation by the dye. Second, while the staining buffer is compatible with other nucleic acid dyes, it is unclear whether it enhances their penetration depth to the same extent as SYTOX Orange. Third, in the 3D histology atlas, some tissues showed some blank space, which was an artefact introduced during cutting.

We have developed COLOR-3D, a simple and scalable method for 3D H&E staining. By providing a reference 3D histology atlas and making discoveries in liver histology, we encourage researchers and histopathologists to apply COLOR-3D to diverse tissue types, accelerating the paradigm shift from 2D histology to 3D histology.

## Methods

### Ethical statement

For mouse tissues, ethics approval was obtained from the Animal Research Ethics Committee of the Chinese University of Hong Kong. Procedures were performed in accordance with the Guide for the Care and Use of Laboratory Animals (AEEC number 20-287-MIS). For post-mortem human tissues, ethics approval was obtained from the Joint Chinese University of Hong Kong-New Territories East Cluster Clinical Research Ethics Committee (approval number 2022.137). For human tumour tissues, ethics approval was obtained from the Joint Chinese University of Hong Kong-New Territories East Cluster Clinical Research Ethics Committee (approval number 2023.407), the Research Ethics Committee (Kowloon Central / Kowloon East) (approval number KC/KE-23-0146/ER-2), and the University of Hong Kong / Hospital Authority Hong Kong West Cluster Institutional Review Board (approval number UW 24-291).

### Chemicals and reagents

SYTOX orange was obtained from Thermo Fisher Scientific (cat. no. S11368). Eosin was obtained from VWR (cat. no. 341972Q). Sulfobutylether-β-cyclodextrin (SBEβCD) was obtained from AraChem (cat. no. CDexB-080/BR). Tetrahydrofuran (THF) was obtained from RCI Labscan. Methanol (MeOH) and dichloromethane (DCM) was obtained from Duksan. Benzyl alcohol and benzyl benzoate were obtained from Sigma Aldrich and Aladdin respectively, and mixed in 1:2 ratio as benzyl alcohol-benzyl benzoate (BABB). Other nucleic acid dyes, lectins and antibodies and their vendors were listed in Supplementary Table 5.

### Human and animal tissues

Adult C478L/6 mice were used. Mice were housed in a controlled environment (22-23°C, 12h light-dark cycle) with unrestricted access to a standard mouse diet and water. The ages of young mice and aged mice were 3-6 months and 36 months respectively. Mouse tissues were perfusion-fixed by 4% paraformaldehyde. Human tissues were fixed for at least 1 week in 10% neutral buffered formalin at room temperature. Porcine liver tissue (for benchmarking experiment and lectin screening) was obtained from a local wet market. Tissues were fixed by 4% paraformaldehyde. All tissue were stored in phosphate buffered saline with 0.02% NaN_3_ (PBSN).

Unless otherwise specified, tissues were processed using a modified iDISCO protocol.^10^ Tissues were dehydrated using 50% MeOH and two changes of 100% MeOH. Tissues were delipidated by immersing overnight in 1:2 MeOH-DCM, then rehydrated with two changes of 100% MeOH. Optionally, tissues were decolorised by RI matching with BABB then illuminated under 40 W LED overnight, followed by two changes of 100% MeOH to remove BABB. The tissues were rehydrated by 50% MeOH, and two changes of PBSN, and stored in PBSN. For all incubation and staining steps, gentle shaking was required. The protocol is summarised in Supplementary Table 6.

### Method development and benchmarking experiments

For method development, pig liver tissues were cut into cubes in 5 mm diameter using a microtome blade. Pig liver tissues were tested against different staining solutions, with a uniform 1-day staining. The basic staining solution contained 7.5 μM SYTOX orange and 36.1 μM eosin in deionised water. Supramolecular hosts including SBEβCD, βCD, 2-hydroxypropyl-β-CD, 2-hydroxypropyl-γ-CD, 4-sulfocalix[4]arene, calix[8]arene, cucurbit[6]uril and cucurbit[7]uril were tested. Supramolecular hosts were added at a concentration of 2.5 mM. Organic solvent including 50% THF, 100% THF, 50% dimethyl sulfoxide (DMSO), 50% ethanol (EtOH), 50% tert-butyl alcohol (t-BuOH), 50% 1-methylimidazole (1-MI), 50% β-phenylethylamine (PEA) and 50% N-butyldiethanolamine (BDEA) were tested. We bisected the cubes and imaged the cut surface using a confocal microscope. The tissue was divided into regions according to distance from the tissue edge, and average staining intensity in each region was measured. Exponential regression was performed. Dye penetration depth was defined as the distance where staining intensity decreases by a factor of 1/e (equation 2). Staining homogeneity was quantified using the coefficient of variation (equation 3).

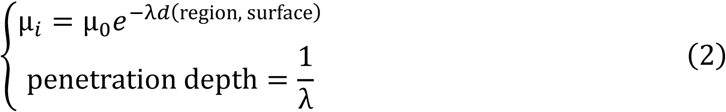

where μ_i_ indicates mean intensity at the i-th region, λ indicates decay constant, and d indicates distance.

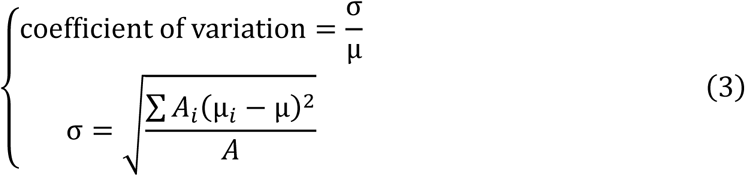

where A_i_ indicates area at the i-th region, A indicates total area, and μ indicates overall mean intensity.

For the benchmarking experiment, we selected state-of-the-art staining protocols from current publications including CUBIC-HV,^16^ CUBIC-HV-500,^16^ Laurino,^13^ and Path3D.^12^ We did not test against CLARITY-based approaches due to their highly specialised protocol with lengthy preprocessing time.^9^ For all protocols, porcine liver tissues were fixed according to the above protocol, but delipidation and staining was performed according to their respective published protocol.

For CUBIC-HV or CUBIC-HV-500, delipidation was performed by immersing in the CUBIC-L solution overnight. Staining was performed by immersing in the ScaleCUBIC-1A or ScaleCUBIC-1A-500 solution respectively. For protocol according to Laurino, delipidation was performed by immersing in 20%, 40%, 60%, 80%, 100% MeOH, where the last step was overnight. Staining was performed by immersing in 100% MeOH. For Path3D, delipidation was performed by immersing in 50% EtOH, two changes of 100% EtOH, followed by xylene overnight. Staining was performed by immersing in Path3D buffer. Due to benchmarking purposes, delipidation and staining duration was standardised across all staining protocols. Staining duration was 24 h across all staining protocols. 7.5 μM SYTOX orange was used across all protocols.

### COLOR-3D protocol

For the complete COLOR-3D protocol, tissues were first fixed and delipidated as described above. COLOR-3D Solution A contained 7.5 μM SYTOX Orange, 2.5 mM SBEβCD, 100 mM NaCl, 50% THF v/v in water. Optionally, tissues were preincubated in COLOR-3D Solution A* for 1 h. Solution A* contained 2.5 mM SBEβCD, 100 mM NaCl, 50% THF v/v in water but without SYTOX Orange. Tissues were then stained in Solution A for 2 days. COLOR-3D Solution B contained 116 μM eosin, 25mM acetic acid / sodium acetate, 95% MeOH v/v in water, adjusted to pH 4. Optionally, tissues were preincubated in Solution B* for 1 h. Solution B* contained 25mM acetic acid / sodium acetate, 95% MeOH v/v in water, adjusted to pH 4 but without eosin. Tissues were then stained in Solution B for 12 h. Tissues were then immersed in 1 change of 100% MeOH and 2 changes of BABB, then imaged with a light sheet or confocal microscope. The above protocol was used for 5-mm thick tissue samples, and the incubation duration (of most steps) was adjusted proportionally. For 1-mm thick tissue samples, the duration was shortened to 1/5 of the original (or 1/4 for rounded numbers). The protocol is summarised in Supplementary Table 6.

For staining of other nucleic acid dyes, the staining solution was obtained by mixing 7.5 μM nucleic dye and COLOR-3D Solution A*. Other steps were identical to the original SYTOX orange protocol. If eosin was not required for staining, the eosin staining step could be skipped from the COLOR-3D protocol.

### Validation and multiplex staining

For immunostaining or lectin staining, we adopted the INSIHGT protocol.^21^ The tissues were first fixed and delipidated as described above. Then, the tissues were pre-incubated in INSIHGT A solution for 1 day, before staining the desired antibody / lectin in INSIHGT A solution for 4 days. All primary antibodies, secondary antibodies and lectin were stained simultaneously. The tissues were then immersed in INSIHGT B solution for 1 day to remove INSIHGT A reagents, followed by two changes of PBSN. After that, COLOR-3D staining was applied to stain SYTOX orange and eosin.

### Liver staining

For liver staining, we used LEL as a liver cell membrane marker. DyLight-594 conjugated LEL was stained in INSIHGT A buffer. SYTOX orange was stained in COLOR-3D Solution A buffer. Eosin was not stained. Details of the protocol were described above.

### Microscopy

For confocal microscopy, we used a Leica SP8 Lightning confocal microscope. Excitation lasers included 405 nm, 488 nm, 514 nm, 552 nm and 638 nm. Detection was used with a 10x, 20x or 40x lens, and a filter with tunable wavelength. For light sheet microscopy, we used a custom built light sheet microscope following the open-source mesoSPIM build plan.^50^ Excitation lasers included 405 nm, 488 nm, 515 nm, 561 nm, 647 nm. Detection was used with Olympus MVX-ZB10 zoom body with a magnification of 0.63x - 6.3x. Filters included QuadLine Rejectionband ZET405/488/561/640, 440/50 ET Bandpass, 509/22 Brightline HC, 542/27 Brightline HC, 585/40 ET Bandpass, 594 LP Edge Basic Longpass, 633 LP Edge Basic Longpass.

### Data processing software

Leica LAS X software was used for viewing image stacks from confocal microscopy. ImageJ and Fiji were used for viewing image stacks from light sheet microscopy as well as processed image stacks.^51^ Matlab 2024b was used to process and analyse image stacks. Python 3.12 and Jupyter Notebook were used to process image stacks and run Cellpose 2.0. Imaris and Imaris Viewer were used to create 3D visualisation and videos of image stacks.

### Image processing and colour conversion pipeline

To process fluorescence image stacks into bright-field H&E, we developed an image processing and colour conversion pipeline. First, we performed image pre-processing including downscaling, 3D Gaussian blur and background subtraction. Second, we performed crosstalk subtraction and intensity scaling by manually examining the optimal crosstalk subtraction and scaling parameters (equation 4). To correct Z-axis intensity attenuation due to photobleaching, an exponential regression was performed to calculate the attenuation factor where each plane was scaled by its reciprocal (equation 5). The image was converted to bright-field H&E (equation 6).^13,20,22^ Thirdly, we performed colour transfer, which mapped the colour distribution of each plane to a reference bright-field H&E image of the same organ. The linear mapping was described by matrix T which matched the mean and covariance matrix of two images (equation 7).^24^ T was obtained from the Linear Monge-Kantorovitch colour transfer equation (equation 8).^24^ Finally, we noticed occasional glitching in the confocal microscope where the motor moved a few optical planes backwards. The glitched planes were detected by a roughness value, followed by thresholding its second derivative. Each glitch was corrected by deleting the planes between the glitched plane and the continuation plane (equation 9)

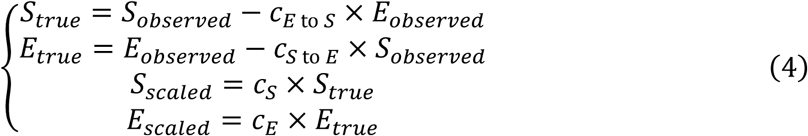

S and E denote the intensity of SYTOX orange and eosin channels respectively. c_E to S_ and c_S to E_ denote the crosstalk coefficients from eosin to SYTOX orange, and vice versa. c_S_ and c_E_ denote the scaling coefficients for SYTOX orange and eosin respectively.

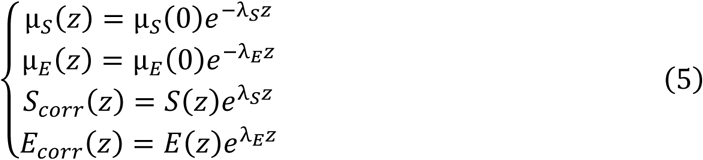

μ_S_(z), λ_S_, S_corr_ denotes mean intensity at z-th plane, decay constant, corrected image for the SYTOX orange channel, and similarly for eosin.

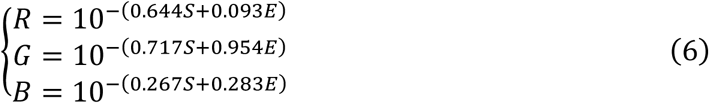

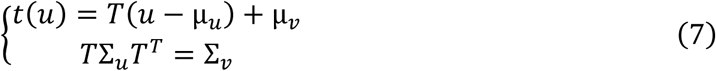

u, v, t(u) and T denote colour vector of original image in RGB, colour vector of reference image, transformed image, and transformation matrix respectively. μ, Σ and T^T^ denote mean of colour distribution, covariance matrix of colour distribution, and transpose of T respectively.

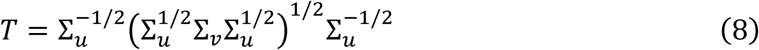

∑^1/2^, ∑^-1/2^ denote positive and negative square roots of ∑.

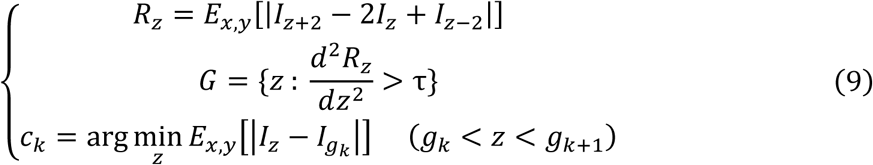

R denotes roughness value. E_x, y_ denotes expected value over the xy plane. I_z_ denotes the z-th plane of the image. G denotes glitched planes. τ denotes threshold. g_k_ and c_k_ denote the k-th glitch plane and continuation plane respectively.

### Liver analysis

Cellpose 2 with custom training was used to segment hepatocyte and hepatocyte nuclei.^34^ Cellpose 2 used a 2.5D approach, which segmented XY, XZ, YZ planes separate and integrated the information. To ensure only hepatocyte and hepatocyte nuclei were segmented, we created a manually segmented dataset from our images and trained custom cell and nuclei segmentation models. We also chose appropriate minimum sizes for hepatocytes and hepatocyte nuclei to filter out segmentation artefacts. We applied morphological opening and closing, and mode filtering to improve mask quality. To assess quality, we extracted random image patches and counted correct segmentation, false positive, false negative and incorrect division. We used Matlab regionprops3 to obtain the following parameters: cell centroid, cell volume, cell eigenvalues, nucleus centroid, nucleus volume and nucleus intensity, and derive N/C ratio and eccentricity (equation 10). We used cell and nucleus centroid to count the number of nuclei and total nuclei intensity in each hepatocyte. We plotted a histogram of total nuclei intensity in each hepatocyte, stratified by cellular polyploidy status. Since there were distinct peaks corresponding to cellular polyploidy status, we set the threshold to be the troughs between two peaks. To ensure the observed nuclei intensity patterns are not due to staining or imaging artefact, we also plotted total nuclear intensity across the x, y, and z axis respectively.

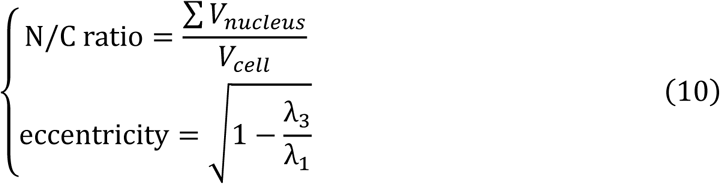

V denotes volume. λ_1_ and λ_3_ denote the largest and smallest eigenvalues.

To segment central veins and portal veins, we exploited the observation that only veins form a large connected empty space in the tissue. We performed inverted thresholding of LEL image, then applied morphological opening and closing. We used bwconncomp to obtain segmentation masks and retained masks with volume larger than 30000 voxels. We labelled each mask as a central vein or portal vein manually since the morphology was readily distinguished. To analyse zonation, we calculated distance ratio based on equation 11.^44^ We separated into 22 zones and discarded the first and last due to the small number of hepatocytes in the first and last zone. Polyploidy status (cellular and nuclear polyploidy) and morphological parameters (cell volume, N/C ratio, eccentricity) were analysed with respect to zonation.

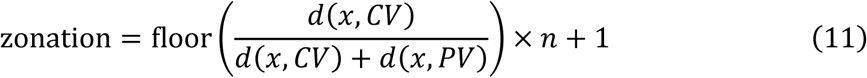

d(x, CV) denotes the distance between the point and nearest central vein. n denotes the number of zones.

For liver lobule shape analysis, we took inspiration from the Voronoi diagram. In a 2D Voronoi diagram, each liver lobule consisted of all points on the 2D plane closest to each CV.^41,42^ The resulting property was that lobule boundaries were equidistant between two CV. Generalisation to 3D faced a limitation since CV is a branching structure in 3D. To solve this challenge, we skeletonised the CV and created a 3D Voronoi diagram where each segment of CV had its own 3D Voronoi region (equation 12). Liver lobule boundaries were defined as Voronoi boundaries between non-adjacent segments. To assess the validity of the Voronoi model, we skeletonised PV and calculated each pixel’s distance from the lobule boundary and the nearest CV respectively (equation 13). Mean and standard deviation of all PV pixels were calculated.

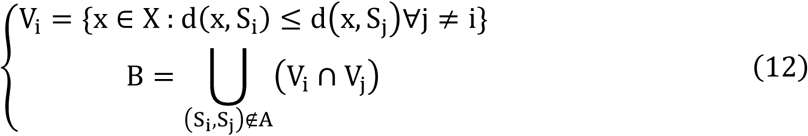

V_i_ denotes the i-th Voronoi region, x denotes any point in the image, X denotes all points in the image, d denotes distance, and S_i_ denotes the CV segment as the generating centre of the i-th Voronoi region. B denotes lobule boundaries, and A denotes the set of adjacent-segment pairs.

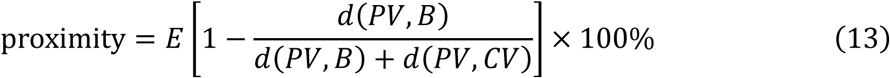

E denotes expected value, and d denotes distance.

### Liver 2D analysis

We performed 2D analysis by reusing the cellular mask, nuclear mask, CV mask and PV mask obtained through 3D analysis. We used the same analysis pipeline except that 3D-based functions were replaced by 2D-based functions, and the threshold of filtering out small objects was adjusted accordingly. For polyploidy analysis, since there were no clear peaks of nuclear intensity even when stratified by cellular polyploidy status, we only analysed cellular polyploidy but not nuclear polyploidy.

### Aged liver analysis

The image analysis pipeline of the aged mouse liver was similar to the young mouse liver. Since aged liver showed varied histology as the young liver, we trained a separate custom model with a dataset from aged mouse liver to ensure accurate segmentation. The image post-processing and regionprops were performed the same as the young mouse. Again, since there were distinct peaks corresponding to cellular polyploidy status, we set the threshold to be the troughs between two peaks. Vessel segmentation was also performed. Since the portal triad was not consistently observed, we did not label veins into central or portal veins, thus zonation analysis and lobule shape analysis were not performed.

### Statistical analysis

The proportion of polyploid hepatocytes were calculated for each liver sample, and confidence interval was calculated based on Student’s t distribution. The Welch’s T test was used to compare the proportions between young and aged liver. No method was used to pre-determine sample sizes.

## Code availability

All codes are available on https://github.com/nickchow246/COLOR-3D-image-analysis/.

## Acknowledgements

We would like to express our deepest gratitude to patients and their families who donated tissues for scientific research. We would like to thank Ms Carmen Chan for administrative assistance in this project. Figures 1A, 1C, 3A, 4A, 5A, 5B, 5F and Supplementary Figure 4 were created with BioRender.com. The project was supported by a Passion for Perfection Program by Faculty of Medicine, The Chinese University of Hong Kong, and a Croucher Innovation Award by The Croucher Foundation, and a Midstream Research Program (MRP/048/20) by the Innovation and Technology Commission.

## Author contributions

Conceptualisation: N.K.N.C., H.M.L. Methodology: N.K.N.C., E.P.L.T., B.T.Y.W., L.Z., J.J.X.L. H.M.L. Investigation: N.K.N.C., E.P.L.T., B.T.Y.W., L.Z., E.T.W.H., Y.T., J.J.X.L., H.M.L. Visualisation: N.K.N.C., H.M.L. Funding acquisition: H.M.L. Project administration: H.M.L. Supervision: H.M.L. Writing: N.K.N.C., H.M.L.

## Competing interests

The authors declare no competing interests.

---

**Supplementary Figure 1:**
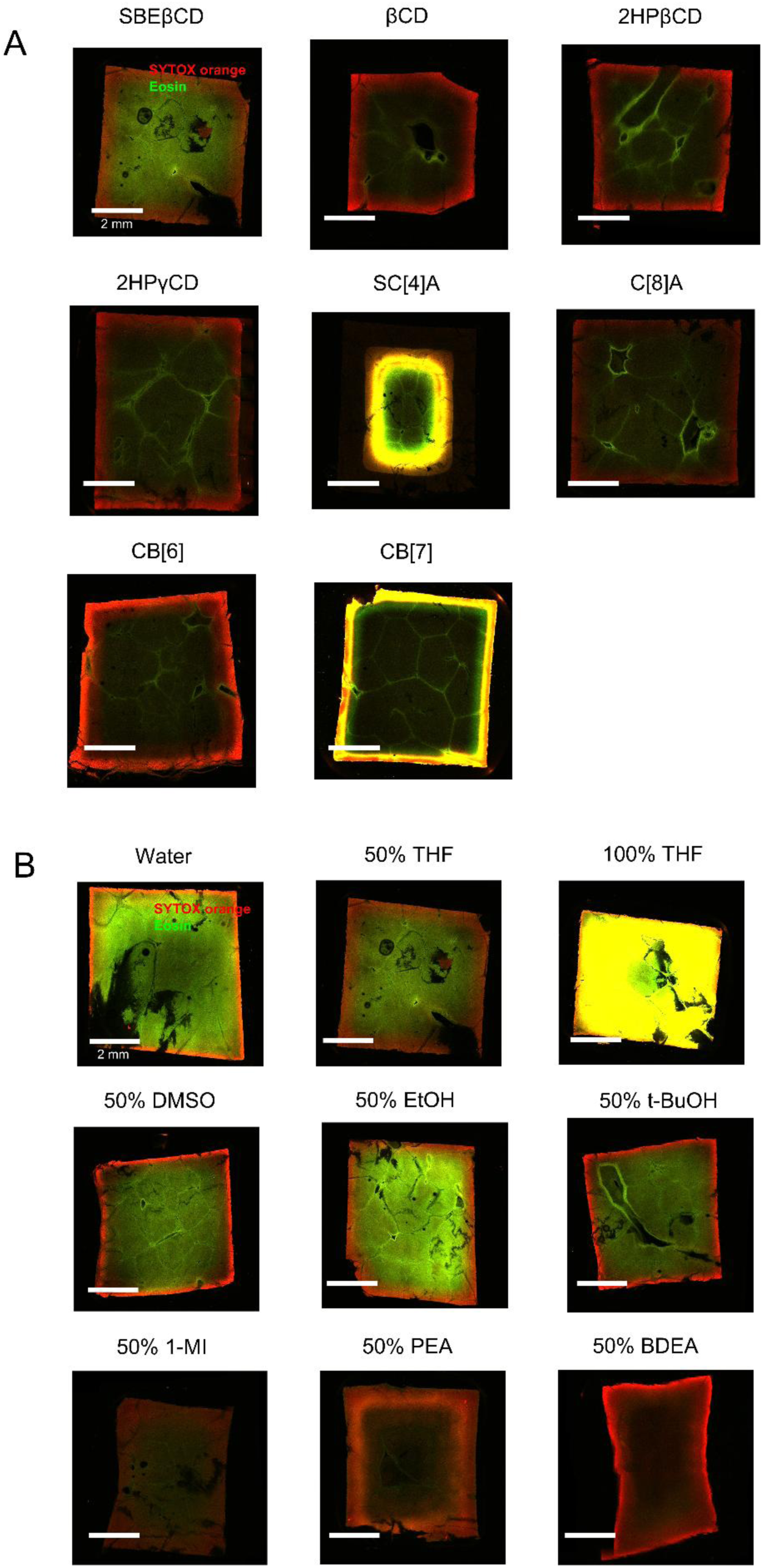
(A) Screening of supramolecular hosts. (2) Screening of organic solvents.

**Supplementary Figure 2:**
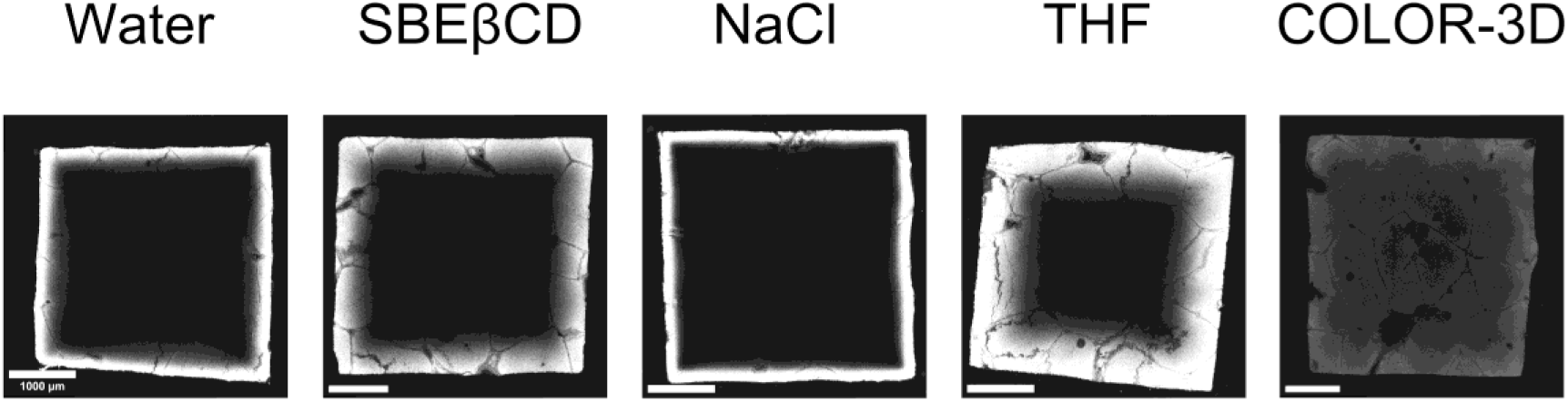
Sulfobutylether-β-cyclodextrin (SBEβCD), tetrahydrofuran (THF) and sodium chloride (NaCl) each improved penetration depth while collectively displaying a synergistic effect.

**Supplementary Figure 3:**
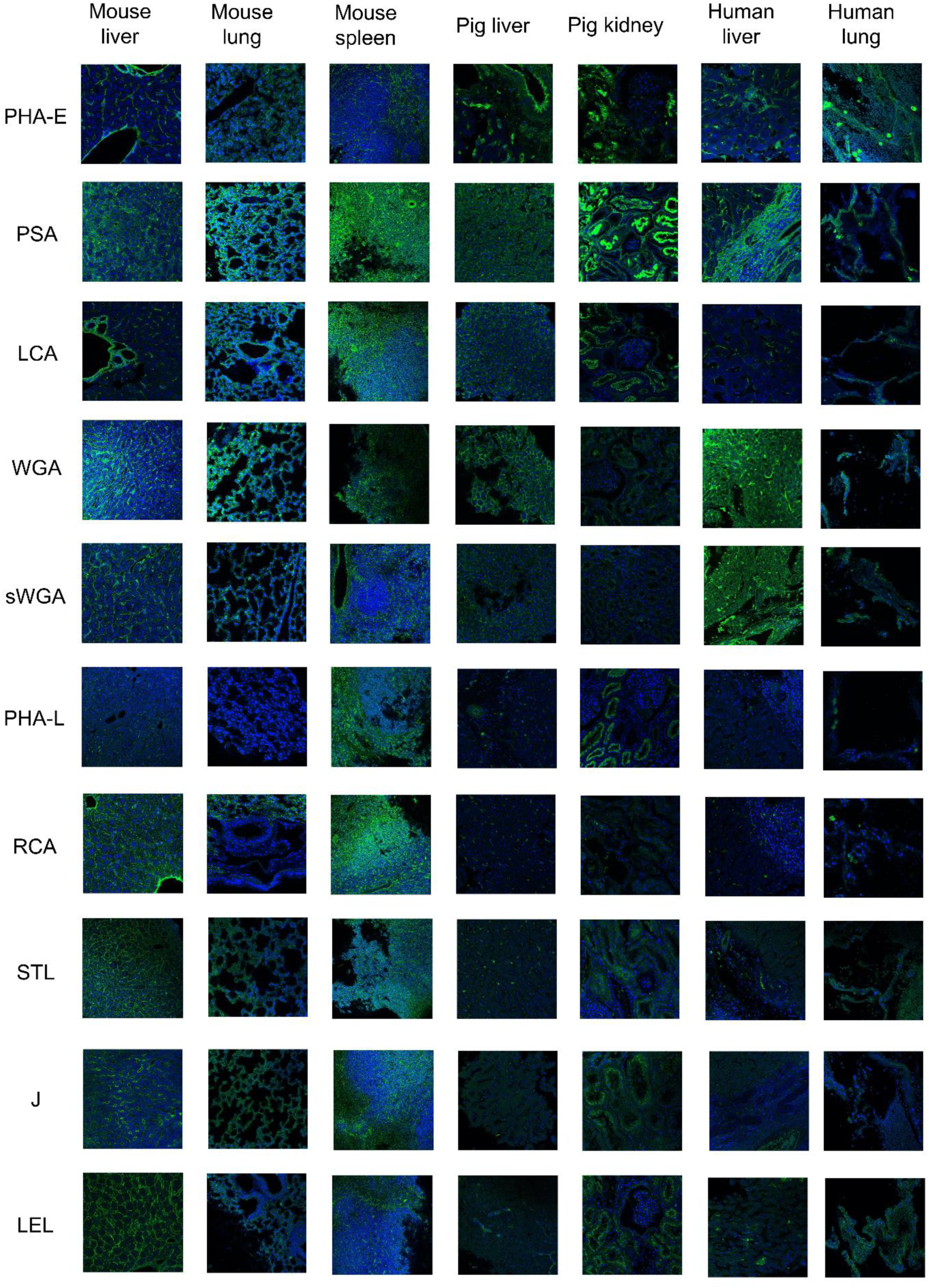
Lectin screening. Lectin is shown as green while DAPI is shown as blue.

**Supplementary Figure 4:**
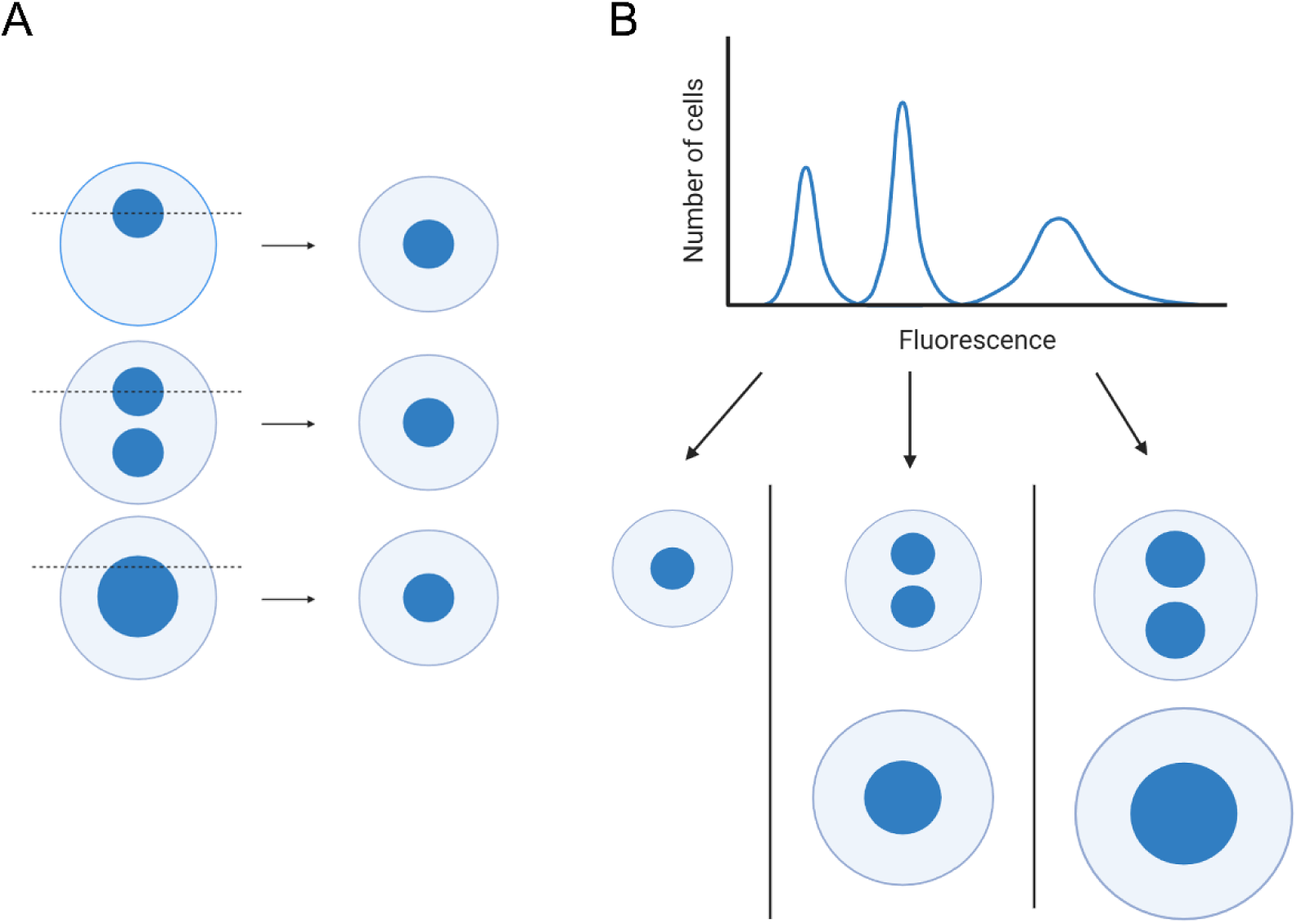
(A) 2D histology underestimates cellular and nuclear polyploidy. (B) Flow cytometry cannot resolve between b2n and m4n hepatocytes, or between b4n and m8n hepatocytes.

**Supplementary Figure 5:**
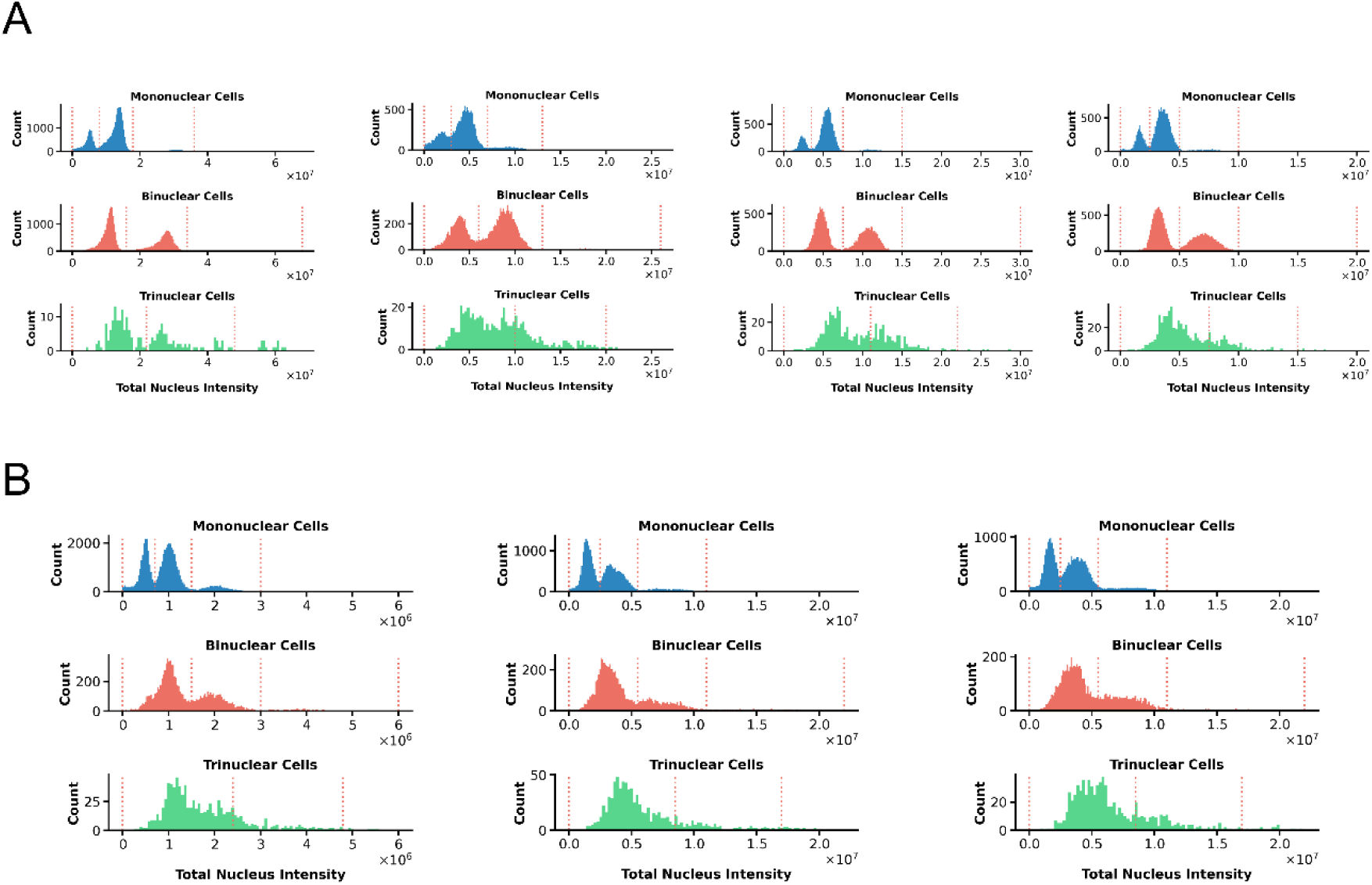
(A) Biological replicates of young mice. (B) Biological replicates of aged mice.

**Supplementary Figure 6:**
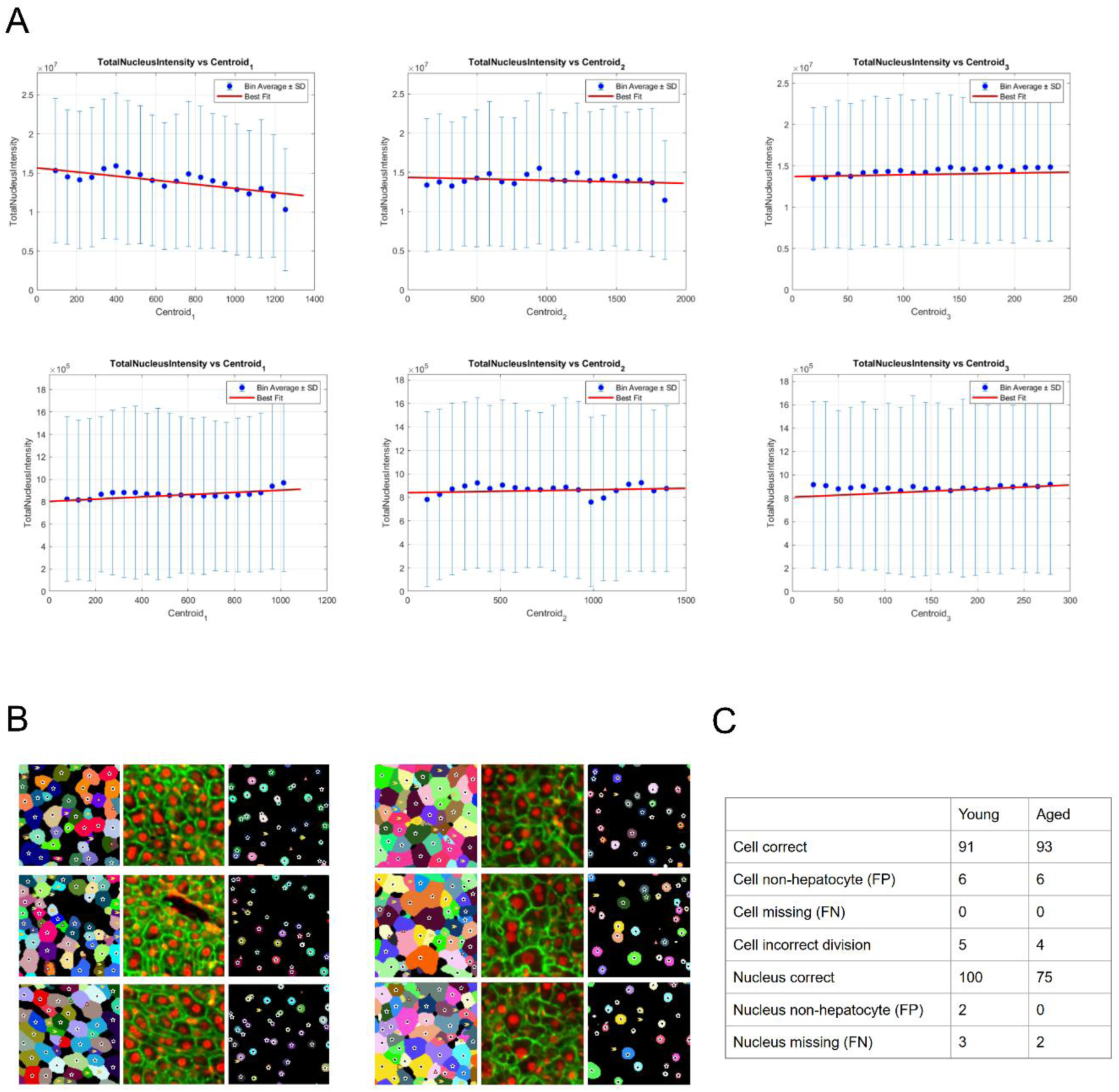
(A) Total nuclear intensity across x, y, z axis for quality control. The top row represents young mice while the bottom row represents aged mice. (B) Segmentation accuracy quality control. Images on the left represent young mice while those on the right represent aged mice.(C) Segmentation accuracy statistics.

**Supplementary Figure 7:**
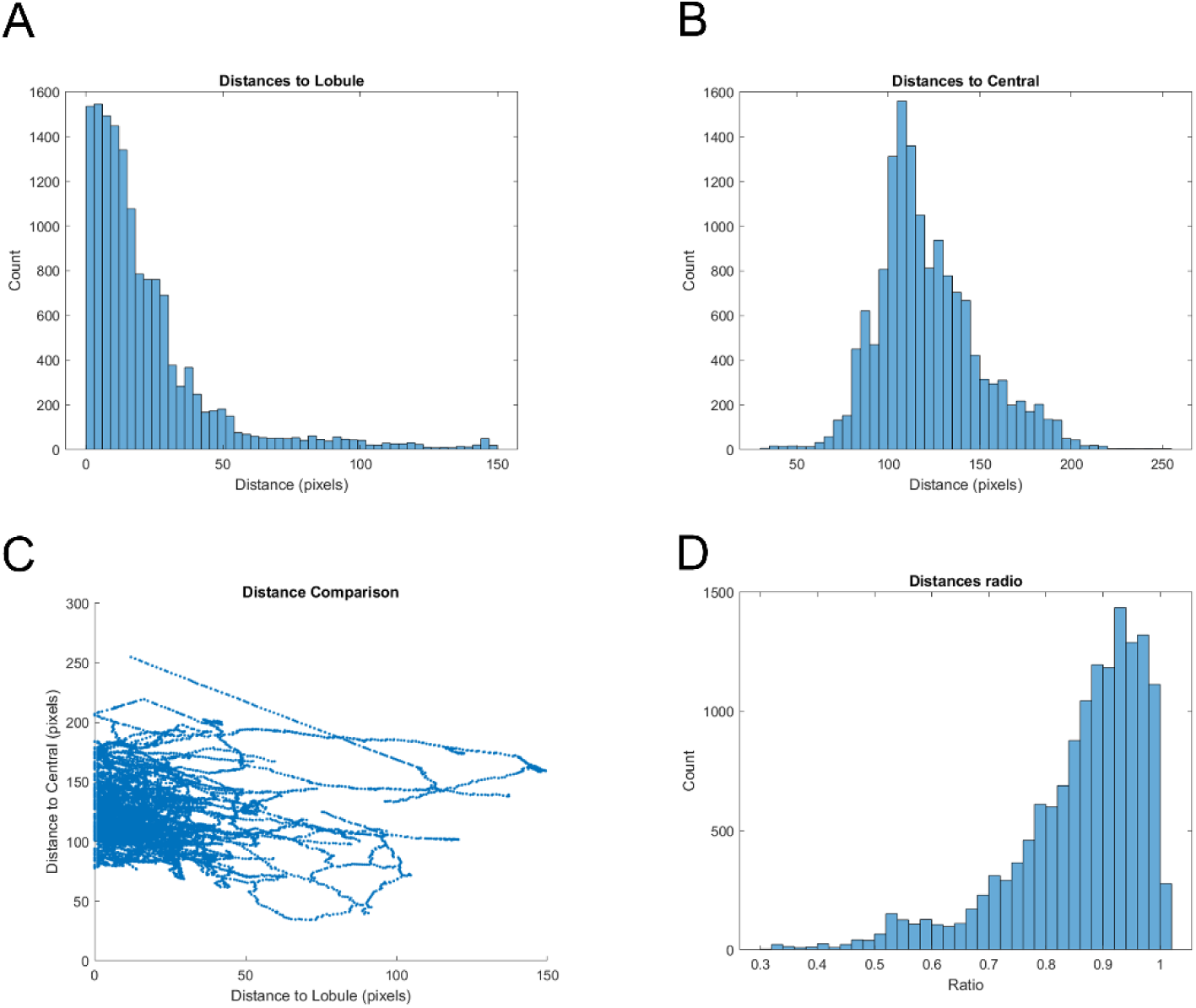
Quantification of the proximity of the portal vein (PV) to the lobule boundary. (A) Distance from the PV to the lobule boundary. (B) Distance from the PV to the central vein (CV). (C) PV-CV distance against PV-lobule boundary distance. (D) Proximity ratio (see Equation 13).

Supplementary Video 1: Mouse ileum in 3D. https://drive.google.com/file/d/1JHWFrQUKLgNd8o-3L_CcU0lZCQ4WZ5xN/view?usp=sharing

Supplementary Video 2: Liver lobule in 3D. https://drive.google.com/file/d/1U6gS227PvMUVLFAPZrCOVhOWjp6sbOcN/view?usp=sharing

Supplementary Video 3: Blood vessels in aged mice liver. https://drive.google.com/file/d/1wohGjCBm01iXSsh_NsRRiOieqPqGAbHF/view?usp=sharing

**Supplementary Table 1:** Representative publications that stain H&E analogues in 3D.

| Protocol name / author / paper (year) | Dye (small molecule dye; others) | Time, tissue (tissue thickness if indicated) | Mechanism of dye penetration |
| --- | --- | --- | --- |
| 3D H&E (light sheet) |  |  |  |
| Bishop (2024) | TO-PRO-3, eosin | 2 days, human prostate punch biopsy | Ethanol-based buffer with NaCl (improve TO-PRO-3 penetration) and weak acid (improve eosin binding) |
| Laurino (2023) | DAPI, SYTOX blue, eosin | 3 days (CLARITY protocol), mouse kidney; 2 days (iDISCO protocol), human bladder (a small piece) | Hydrogel (CLARITY protocol); treatment with MeOH (iDISCO protocol) |
| Freise (2023) | Acridine orange, eosin | 1 day, human placenta (1.5 mm) | Triton X-100 (permeabilisation) |
| Li (2020) | TO-PRO-3, eosin | 2 days, human lung (0.9 mm) | Triton X-100 (permeabilisation) |
| Susaki (2020) | Propidium iodide, SYTO 16, SYTOX Green, BOBO-1, RedDot2, eosin | 5 days, mouse brain | Triton X-100 (permeabilisation); Quadrol and urea (reduced dye / tissue interaction) |
| 3D H&E (confocal / two-photon) |  |  |  |
| Peng (2022) | SYTO 16, DiD | 4 h, human prostate (100 µm) | Triton X-100 (permeabilisation) |
| Kim (2022) | SYTO 16, eosin | 1 day, mouse kidney / human brain / etc. (1 mm) | Hydrogel (modified CLARITY protocol) |
| Giacomelli (2016) | DAPI, eosin | Human breast (40 µm) | None |
| Torres (2014) | SYTOX Green, acridine orange | 2 days; human prostate / liver / etc. (1 mm) | None |

**Supplementary Table 2:** Morphological description of notable features observed in 3D.

| Organ | Structure / morphology |
| --- | --- |
| Human lung | Size / difference in size of alveoli<br>Connection between alveoli |
| Human colon gland | Colonic gland and lamina propria |
| Human colon cancer | Cribriform gland<br>Eosinophil and lymphocyte infiltration<br>Tumour cells encapsulating submucosal plexus |
| Mouse tongue epithelium | Filiform papillae, connected with muscle layer |
| Mouse tongue muscularis externa | Orientation of muscle fibre<br>Nerve, vessel |
| Mouse cecum | Relationship between villi / crypt / lamina propria |
| Mouse ileum | Relationship between villi / crypt / lamina propria<br>Myenteric ganglion<br>Inner and outer smooth muscle direction |
| Mouse thymus | Hassall corpuscle |
| Mouse lung | Size / difference in size of alveoli<br>Connection between alveoli |
| Mouse pancreas | Acini, duct, islet<br>1-cell-thick Intercalated duct clearly shown in 3D |
| Mouse spleen | White pulp and red pulp<br>Splenic sinus clearly shown in 3D |
| Human leiomyosarcoma | Smooth muscle bundles have random orientation<br>Frequent mitosis and atypical mitosis clearly shown in 3D |
| Mouse heart | Cardiomyocyte and purkinje fiber<br>Purkinje-cardiomyocyte contact |
| Human breast cancer | Invasive duct with occasional single-file pattern<br>3D helps clarify if single-file pattern is cross-section of a gland |
| Human brain | Neuron, glial cell, neuropil |

**Supplementary Table 3:** Comparison of the proportion of polyploid hepatocytes between young and aged mice.

|  | young_m<br>ean | young_sd | young_ci<br>_margin | aged_me<br>an | aged_sd | aged_ci_<br>margin | t_stat | p_value | stars |
| --- | --- | --- | --- | --- | --- | --- | --- | --- | --- |
| Mononuc<br>lear<br>(Total) | 47.52 | 2.65 | 4.21 | 78.84 | 3.43 | 8.51 | -13.16 | 0.0003 | *** |
| 2n<br>(Mono) | 11.36 | 2.18 | 3.46 | 32.37 | 3.5 | 8.69 | -9.16 | 0.0023 | ** |
| 4n<br>(Mono) | 33.43 | 1.3 | 2.07 | 37.87 | 5.26 | 13.07 | -1.43 | 0.2791 | ns |
| 8n<br>(Mono) | 2.64 | 0.7853 | 1.25 | 8 | 1.3 | 3.23 | -6.34 | 0.0072 | ** |
| >8n<br>(Mono) | 0.094 | 0.0391 | 0.0622 | 0.6005 | 0.0656 | 0.163 | -11.88 | 0.0012 | ** |
| Binuclear<br>(Total) | 50.41 | 2.02 | 3.22 | 18.74 | 2.69 | 6.68 | 17.09 | 0.0001 | *** |
| 2n (Bi) | 26.18 | 5.14 | 8.19 | 12.47 | 2.54 | 6.3 | 4.63 | 0.0072 | ** |
| 4n (Bi) | 23.64 | 3.28 | 5.22 | 5.38 | 0.5432 | 1.35 | 10.93 | 0.0012 | ** |
| 8n (Bi) | 0.5979 | 0.3343 | 0.5319 | 0.8632 | 0.0963 | 0.2392 | -1.51 | 0.2134 | ns |
| >8n<br>(Bi) | 0.0004 | 0.0008 | 0.0013 | 0.0303 | 0.0106 | 0.0264 | -4.86 | 0.0392 | * |
| Trinuclea<br>r (Total) | 1.6 | 0.8457 | 1.35 | 2.13 | 0.6324 | 1.57 | -0.9505 | 0.3857 | ns |
| 2n (Tri) | 1.14 | 0.6486 | 1.03 | 1.67 | 0.47 | 1.17 | -1.25 | 0.2683 | ns |
| 4n (Tri) | 0.4402 | 0.2396 | 0.3813 | 0.4002 | 0.1478 | 0.3673 | 0.2716 | 0.797 | ns |
| >4n<br>(Tri) | 0.0181 | 0.0049 | 0.0079 | 0.0625 | 0.0386 | 0.0958 | -1.98 | 0.183 | ns |
| Tetranucl<br>ear<br>(Total) | 0.4071 | 0.3097 | 0.4929 | 0.2483 | 0.1091 | 0.271 | 0.9499 | 0.397 | ns |

**Supplementary Table 4:** Comparison on the proportion of 2n, 4n, 8n and 16n nuclei between young and aged mice.

|  | young_mean | young_sd | young_ci_margin | aged_mean | aged_sd | aged_ci_margin | t_stat | p_value | stars |
| --- | --- | --- | --- | --- | --- | --- | --- | --- | --- |
| 2n | 44.4 | 5.45 | 8.68 | 51.04 | 6.54 | 16.24 | -1.43 | 0.2282 | ns |
| 4n | 53.02 | 4.65 | 7.4 | 40.39 | 5.46 | 13.57 | 3.22 | 0.0324 | * |
| 8n | 2.52 | 0.9556 | 1.52 | 8.04 | 1.07 | 2.67 | -7.05 | 0.0019 | ** |
| 16n | 0.0617 | 0.0267 | 0.0426 | 0.5338 | 0.0599 | 0.1489 | -12.73 | 0.002 | ** |

**Supplementary Table 5:** List of antibodies, lectins and nucleic acid dyes.

| Antibody / lectin / nucleic acid dye | Vendor | Catalogue number |
| --- | --- | --- |
| ABCC2 | Abclonal | A8405 |
| KRT7 | Abclonal | A2574 |
| KRT8 | Abclonal | A1024 |
| GAP43 | Novus Bio | NB300-143 |
| Phaseolus vulgaris erythroagglutinin (PHA-E) | Vector Laboratories | RL-1122-2 |
| Pisum sativum agglutinin (PSA) | Vector Laboratories | RL-1052-5 |
| Lens culinaris agglutinin (LCA) | Vector Laboratories | RL-1042-5 |
| Wheat germ agglutinin (WGA) | Vector Laboratories | RL-1022-5 |
| Succinyl wheat germ agglutinin (sWGA) | Vector Laboratories | RL-1022S-5 |
| Phaseolus vulgaris leucoagglutinin (PHA-L) | Vector Laboratories | RL-1112-2 |
| Ricinus communis agglutinin (RCA) | Vector Laboratories | FL-1081-1 |
| Solanum tuberosum lectin (STL) | Vector Laboratories | FL-1161-2 |
| Jacalin (J) | Vector Laboratories | FL-1151-5 |
| Lycopersicon esculentum lectin (LEL) | Vector Laboratories | DL-1178-1 |
| Griffonia simplicifolia lectin 1 (GSL1) | Vector Laboratories | DL-1208-5 |
| DAPI | Thermo Fisher Scientific | D1306 |
| Syto 16 | Thermo Fisher Scientific | S7578 |
| Yoyo 3 | Thermo Fisher Scientific | Y3606 |

**Supplementary Table 6:** Summary of the COLOR-3D protocol.

| # | Chemical | Volume<br>(compared to<br>tissue sample) | Time (1 mm) | Time (5 mm) |
| --- | --- | --- | --- | --- |
| Fixation |  |  |  |  |
| 1 | 4% paraformaldehyde | 10x | 1 day | 1 day |
| 2 | Phosphate-buffered saline with 0.02% NaN<sub>3</sub> (PBSN) | 10x | Until use | Until use |
| Delipidation |  |  |  |  |
| 3 | 50% methanol (MeOH) | 5x | 15 min | 1 h |
| 4 | 100% MeOH x 2 | 5x | 15 min x 2 | 1 h x 2 |
| 5 | 1:2 MeOH-DCM | 5x | 1 day | 1 day |
| 6 | 100% MeOH x 2 | 5x | 15 min x 2 | 1 h x 2 |
|  | Optional steps: 1:2 benzyl alcohol-benzyl benzoate (BABB) → LED overnight → 100% MeOH x 2 |  |  |  |
| 7 | 50% MeOH | 5x | 15 min | 1 h |
| 8 | PBSN x 2 | 5x | 15 min x 2 | 1 h x 2 |
| Immunostaining with INSIHGT (if applicable) |  |  |  |  |
| 9 | INSIHGT A (pre-incubation) | 1x | 6 h | 1 day |
| 10 | INSIHGT A (immunostaining) | 1x | 1 day | 4 days |
| 11 | INSIHGT B | 3x | 6 h | 1 day |
| 12 | PBSN x 2 | 5x | 15 min x 2 | 1 h x 2 |
| COLOR-3D staining |  |  |  |  |
| 13 | COLOR-3D Solution A* | 5x | 15 min | 1 h |
| 14 | COLOR-3D Solution A | 1x | 12 h | 2 days |
| 15 | COLOR-3D Solution B* | 5x | 15 min | 1 h |
| 16 | COLOR-3D Solution B | 1x | 3 h | 12 h |
| Refractive index matching |  |  |  |  |
| 17 | 100% MeOH | 5x | 15 min | 1 h |
| 18 | Benzyl alcohol benzyl-benzoate (BABB) x 2 | 5x | 30 min x 2 | 2 h x 2 |

## Notes

### Competing Interest Statement

The authors have declared no competing interest.

